# Quantitatively Monitoring *in situ* Mitochondrial Thermal Dynamics by Upconversion Nanoparticles

**DOI:** 10.1101/2020.11.29.402818

**Authors:** Xiangjun Di, Dejiang Wang, Jiajia Zhou, Lin Zhang, Martina Stenzel, Qian Peter Su, Dayong Jin

## Abstract

Temperature dynamics reflect the physiological conditions of cells and organisms. Mitochondria regulates temperature dynamics in living cells, as they oxidize the respiratory substrates and synthesize ATP, with heat being released as a by-product of active metabolism. Here, we report an upconversion nanoparticles based thermometer that allows *in situ* thermal dynamics monitoring of mitochondria in living cells. We demonstrate that the upconversion nanothermometers can efficiently target mitochondria and the temperature responsive feature is independent of probe concentration and medium conditions. The relative sensing sensitivity of 3.2% K^−1^ in HeLa cells allows us to measure the mitochondrial temperature difference through the stimulations of high glucose, lipid, Ca^2+^ shock and the inhibitor of oxidative phosphorylation. Moreover, cells display distinct response time and thermal dynamic profiles under different stimulations, which highlights the potential applications of this thermometer to study *in situ* vital processes related to mitochondrial metabolism pathways and interactions between organelles.

## Introduction

Intracellular temperature is a crucial parameter to assess the status of living cells and organisms ^1^. The activations of a wide range of chemical reactions in the living cell, especially in the mitochondria, produce a large amount of energy and cause the change of temperature. As the powerhouse of the cell, mitochondria provide energy to the living cell through the oxidative phosphorylation process ^2^. During this chemical reaction, about 67% of the energy is used to synthesize ATP and the other ~33% dissipates in the form of heat ^3^. Failure to produce ATP will cause a change of mitochondria temperature, so that the variation of mitochondria temperature indicates the cellular metabolism status ^4^. Given its importance to fundamental studies, disease diagnosis, and therapy, accurate and specific temperature sensing remains as a challenge due to the lack of noninvasive sensing probes ^5–8^.

Fluorescence nanothermometry has emerged to noninvasively reveal the localized intracellular temperature in living cells ^9^. Temperature responsive fluorescent materials ^10–12^, including small molecules ^13^, fluorescent polymers ^1^, fluorescent proteins ^14^, and inorganic particles ^15, 16^, have been extensively explored. For example, Homma *et al.* developed a ratiometric nano-thermosensor (Mito-RTP) by using thermos-sensitive rhodamine B and thermo-insensitive CS NIR dye that enable the temperature monitoring of mitochondria under chemical stimulation ^13^. Yang *et al.* designed photoluminescence spectral shifts quantum dots (QDots) to monitor temperature change in NIH/3T3 cells under Ca^2+^ stress and cold shock ^15^. However, due to the photo-bleaching and photo-blinking properties, these fluorescent nanothermometers limit in the area of long-term tracking and sensing. Lanthanide doped upconversion nanoparticles (UCNPs) with stable photonic characteristics are suitable for long-term bio-sensing, bio-imaging ^17, 18^, and photothermal therapy ^19^. UCNPs are chemically stable ^17^, optically stable ^20^, and biologically compatible ^21^. The anti-Stokes emission process based on NIR light excitation avoids cellular auto-fluorescence and photo-damage but has a high tissue penetration ^21^.

The first demonstration of UCNPs nanothermometers has successfully monitored around 5 °C temperature change of living cells upon external heating in 2010 ^20^. The UCNPs doped with erbium ions demonstrate a temperature-dependent luminescence following the Boltzmann distribution ^22^,

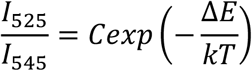

where *I*_*525*_ and *I*_*545*_ are the integrated fluorescent intensities around 525 nm and 545 nm emission peak, respectively; *C* is a constant; *ΔE* is the energy gap; *k* is the Boltzmann constant and *T* is the absolute temperature. Recently, the UCNPs nanothermometers have been also applied for *in vivo* small animal imaging and temperature monitoring ^17, 23^. Shi *et al.* used UCNPs doped with Nd^3+^ to sense temperature changes in NIH-3T3 cells ^17^. Qiu *et al.* fabricated a hybrid structure composed of PbS quantum dots and Tm- doped UCNPs to realize intratumoral monitor *in vivo* ^23^. However, there remains a big gap to realize the UCNPs thermometry with *in situ* organelle targeting capability for localized intracellular temperature sensing. It is challenging to modify them into biocompatible and specific labeling through incubation. This is because, for relatively large inorganic nanoparticles (e.g., 20 nm) with native positive charges from lanthanide ions.

In this study, by using mitochondria specifically, temperature-dependent and non-photobleaching UCNPs, we aimed to monitor the *in situ* mitochondrial temperature dynamics specifically under different nutrient conditions and chemical stimulations. A crosslinked polymer network was applied to avoid the aggregation of UCNPs in the buffer ^24^. Then PEGMEMA_80_-*b*-EGMP_3_ di-block copolymers and 4Arm-PEG-NH_2_ were conjugated to the UCNPs to make them dispersible in buffer and cell culture medium with functional amine groups on the surface. Lastly, (3-carboxypropyl)triphenylphosphonium bromide (TPP), as a targeting moiety, was covalently bond to UCNPs. This strategy leads to the nanoparticles capable of targeting mitochondria ^25, 26^. The practicality of the temperature-sensing was confirmed by real-time monitoring of the mitochondrial temperature variations induced by external nutrient conditions and chemical stimulations, including glucose, lipid, Ca^2+^, and the inhibitor of oxidative phosphorylation. With the nanothermometer, mitochondria display different profiles of reaction time and thermal dynamics under different physiological nutrient conditions and chemical stimulations. Interestingly, mitochondria respond faster and stay longer at a relatively high-temperature level in high oleic acid versus high glucose culture medium, which indicates different pathways of glycometabolism and lipid metabolism. The distinct thermodynamics highlights the extensive applications of this thermometer to study vital biological processes related to mitochondrial metabolism pathways and interactions between mitochondria and other organelles, like lysosome, ER ^27^ Golgi ^28^, lipid droplet, and peroxisome ^29^.

## Results

### Design of UCNPs@TPP

The method to synthesize UCNPs was described previously (see details in the Methods section) ^30^. To construct a stable and dispersible mitochondria targeting thermosensor, a crosslinked polymer network was applied to modify the UCNPs surface (**Figure 1A**) ^24^. These hydrophilic, crosslinked coating layers are hard to detach from the particles and keep UCNPs stable in the culture media and the intracellular environment (see Methods for detailed reactions). The transmission electron microscopy (TEM) images (**Figure 1B-E**) showed the morphology uniformity and monodispersity of the nanoparticles before and after the modification. The dynamic light scattering (DLS) results (**Figure 1F**) showed the high uniformity with the hydrodynamic size increasing from 31.09 ± 2.76 nm of the as-synthesized UCNPs to 39.43 ±1.64, 42.34 ± 2.54, and 45.13 ± 2.55 nm (measured by TEM in **Figure 1B**). For TPP conjugation, the amine groups of 4Arm-PEG-NH_2_ on the crosslinked polymer network provide the anchoring groups for TPP. The Zeta potential results in **Figure 1G** indicated the successful modification of each step, as the surface charge turns from negative 13 mV to positive 18 mV with the exposure of 4Arm-PEG-NH_2_ and positive 32 mV with TPP by a carbodiimide reaction ^31, 32^. The strong positive charge on the UCNPs surface facilitated the nanoparticles to locate into the cell cytoplasm and arrive at the mitochondria ^26^. Furthermore, the long-term stability of UCNPs@TPP was tested by DLS (**Figure S1A**). These particles all exhibited excellent dispersity and stability in the complete medium (Dulbecco’s Modified Eagle’s Medium (DMEM) containing 10% v/v fetal bovine serum (FBS) and 1% v/v penicillin-streptomycin) and the incubation medium (DMEM containing 2% FBS and 0.5% v/v BSA), as shown in **Figure S1B**.

**Figure 1.**
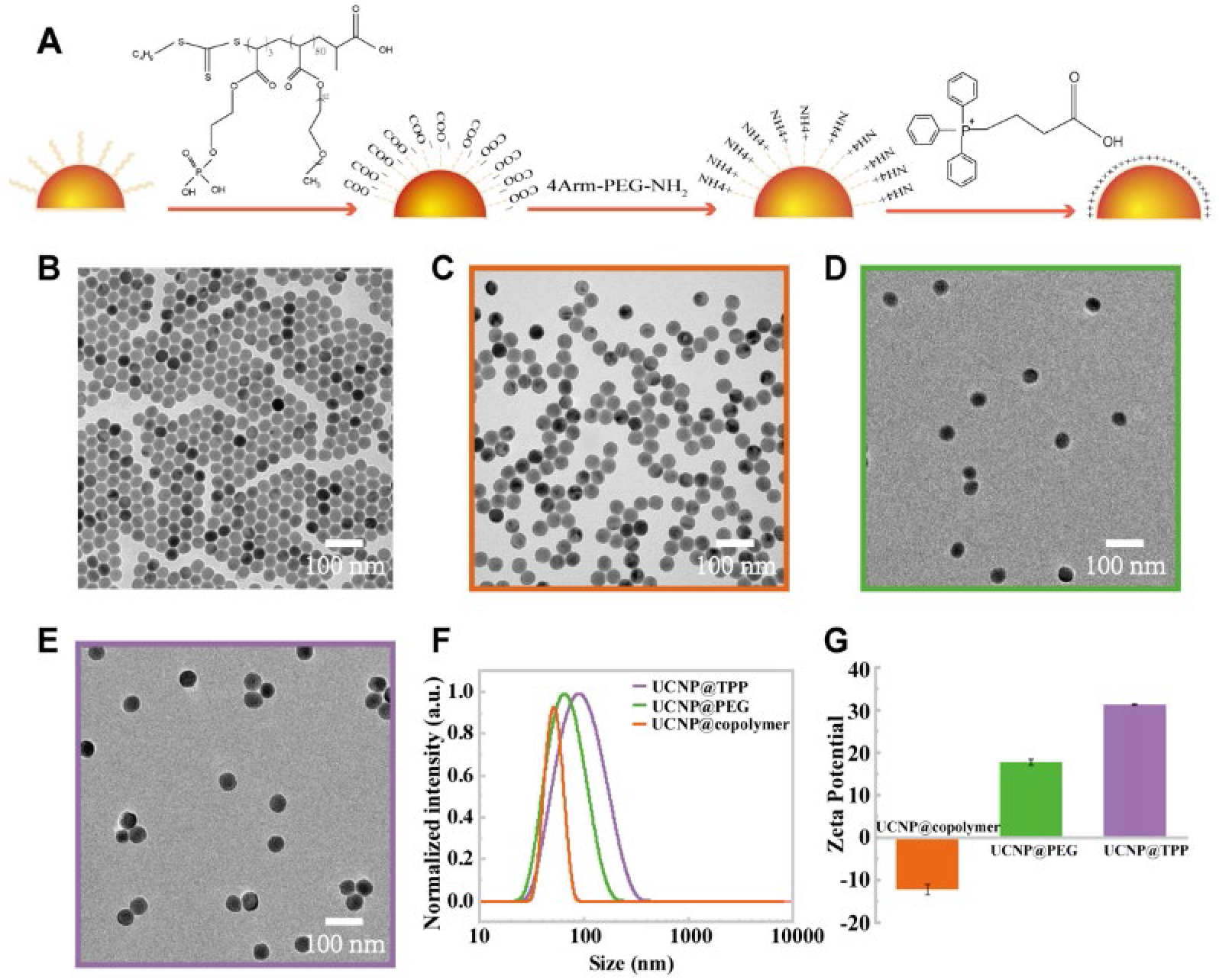
Characterization of UCNPs thermosensors. A. Schematic illustration of the mitochondria-targeted probes with crosslinked polymer layers and TPP. B-E. Representative TEM images of UCNPs, UCNPs@copolymer, UCNPs@PEG, and UCNPs@TPP. F. DLS for UCNPs with different surface modifications. G. Zeta potential of UCNPs with different surface modifications.

### Thermoresponsive properties of UCNPs@TPP *in vitro*

Under the 980 nm excitation, the modified nanoparticles UCNPs@TPP emitted green emission consists of two distinct bands between 515-535 nm (centered at 525 nm) and 535-570 nm (centered at 545 nm) (**Figure 2A**), attributing to the ^2^H_11/2_ and ^4^S_3/2_ transitions of Er^3+^, respectively. While the emitted intensities at both 525 nm and 545 nm peaks decreased when the temperature elevated from 30 °C to 60 °C due to thermal quenching, the ratio of 525 nm and 545 nm peaks increased by following the law of Boltzmann distribution (**Figure 2B**).

**Figure 2.**
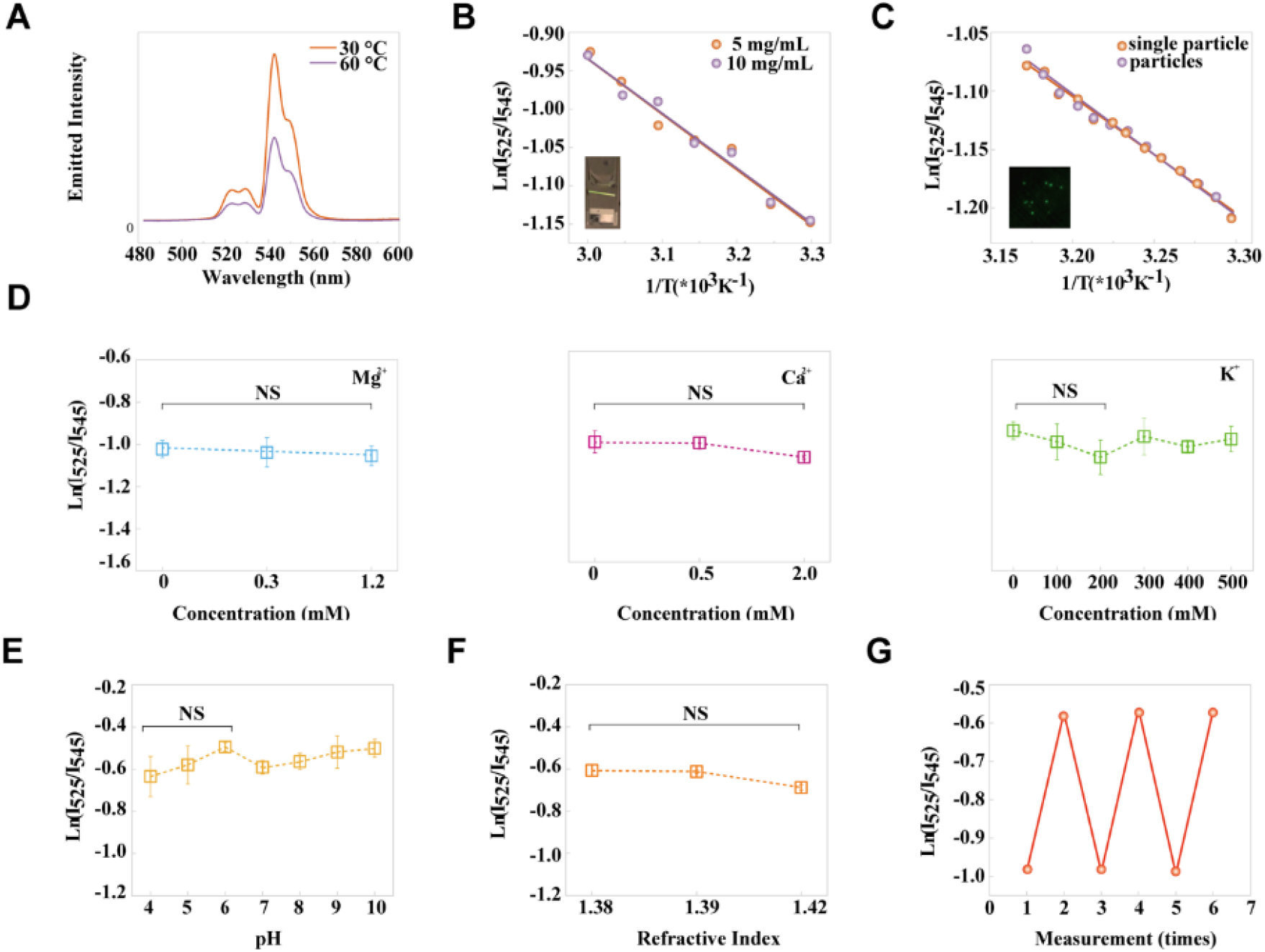
Thermoresponsive properties of UCNPs@TPP in vitro. A. Upconversion emission spectra obtained at two different cuvette temperatures (λexc = 980 nm). B. Plots of Ln(I_525_/I_545_) vs 1/T to calibrate the thermometric scale in the solution with different concentrations. C. Plots of Ln(I_525_/I_545_) vs 1/T to calibrate the thermometric scale at the single-particle level versus multiple particle level. D. Thermal sensitivity of UCNPs@TPP in response to Mg^2+^(left), Ca^2+^(middle) or K^+^(right) (n = 3 independent experiments). E. Changes in the Ln(I_525_/I_545_) ratio of UCNPs@TPP in response to pH (n = 3 independent experiments). F. Changes in the Ln(I_525_/I_545_) ratio of UCNPs@TPP in response to refractive index (n = 3 independent experiments). G. Reversibility of the temperature-dependent changes of UCNPs@TPP fluorescence. The solution temperature was changed from 30 °C to 45 °C. Data points represent mean ± s.d.

An accurate calibration curve is extremely crucial to quantitatively analyze temperature dynamics in the intracellular environment. Ideal fluorescent thermosensors should be independent of the concentration as it’s difficult to control and measure the concentration of fluorescent thermosensors in living cells. As shown in **Figure 2B**, by calibrating two solutions containing different concentrations of UCNPs@TPP (5 and 10 mg/mL) using a purpose-built spectrometer, the Residual Sum of Squares (RSS) of these two linear fittings was measured as 0.013159, which indicates that UCNPs@TPP as a fluorescent thermosensor is concentration-independent. To confirmed this result, the calibration curves at the single-particle level were performed with a purpose-built total internal reflected fluorescent (TIRF) microscopy system. Although the calibration obtained by microscope had distinction from those by spectrometer because of the different sensitivities of the spectrometer with photomultiplier tube (PMT) and microscope with Electron-multiplying CCD (EMCCD), the RSS of these two linear fittings was measured as 0.00482. Notably, the thermal responsiveness of UCNPs@TPP was essentially unchanged under different concentrations whether from single particle level or in the solution.

The micro-environment conditions in living cells, like the ionic strength, pH value, and refractive index vary as time and locations change, also vary from one organelle to another. For example, the refractive indexes of the cell cytoplasm, nucleus, and mitochondria are 1.38, 1.39, and 1.42 respectively ^33^. Fluorescent thermosensors should keep stable in the living cell. **Figure 2D-F** illustrates that temperature-dependent fluorescence changes in UCNPs@TPP were unaffected by Mg^2+^ or Ca^2+^ at different physiological intracellular concentrations, ionic strength (0-500 mM), pH (4-10), and refractive indexes (1.38-1.42). Furthermore, the I_525_/I_545_ ratios of UCNPs@TPP can be reproduced by heating and cooling between 30 °C and 45 °C for three cycles, demonstrating the reversibility of UCNPs@TPP without thermal denaturation.

Since the absorption spectrum of water is in the range of 680-1000 nm with the peak at 980 nm, we monitored the temperature variations of UCNPs@TPP excited by a 980 nm laser at an intensity of 0.5 kW/cm^2^ for 30 minutes. As shown in **Figure S2**, an illumination of 0.5 kW/cm^2^ 980 nm laser did not lead to temperature elevation of the water within 30 minutes.

### UCNPs@TPP work as subcellular thermosensors in HeLa cells

To apply UCNPs@TPP in live cells, we first checked the cytotoxicity of UCNPs@TPP in live cells with short-time treatment at different concentrations measured by MTT test experiments. **Figure S3** showed that the cell viability in the experimental groups was similar to that of the control group, indicating that UCNPs@TPP has negligible cytotoxicity to HeLa cells. Considering the labeling efficiency, 50 μg/mL was chosen in the following live-cell experiments.

Next, the exact locations of nanoparticles in the live cell were tested. Colocalizations of UCNPs@TPP and MitoTracker Deep Red were conducted by a purpose-built TIRF microscope with 980 nm and 647 nm lasers as the light sources. As shown in **Figure 3A**, the green channel illustrated that these three kinds of nanoparticles dispersed into HeLa cells after 12 hours’ incubation. UCNPs@copolymer and UCNPs@PEG preferred to aggregate, while UCNPs@TPP dispersed well within the cell. The merged images showed UCNPs@TPP had a better colocalization than those of control groups. Pearson’s R-value for the experimental group was 0.70. In contrast, Pearson’s R-value for the UCNPs@PEG and UCNPs@copolymer treatment groups were 0.40 and 0.27, respectively. These results indicated that UCNPs@TPP preferred to target the mitochondria. As the focus, the temporal and spatial resolutions of the microscope may affect the accuracy of colocalization results, we further confirmed the locations of the particles by isolating the mitochondria from HeLa cells using a mitochondria isolation kit (Thermo Fisher), dispersed in 50 μL of PBS buffer. The mitochondria suspension was transferred to a 96-well plate and dried at room temperature. Then the fluorescence intensity of the isolated mitochondria was measured. In the control groups (**Figure 3B)** with UCNPs@PEG and UCNPs@copolymer, the fluorescence intensities of nanoparticles were barely seen with the mitochondria, while the intensity of UCNPs@TPP treated group was ~3 times higher than those of the other two groups, indicating the successful mitochondria targeting.

**Figure 3.**
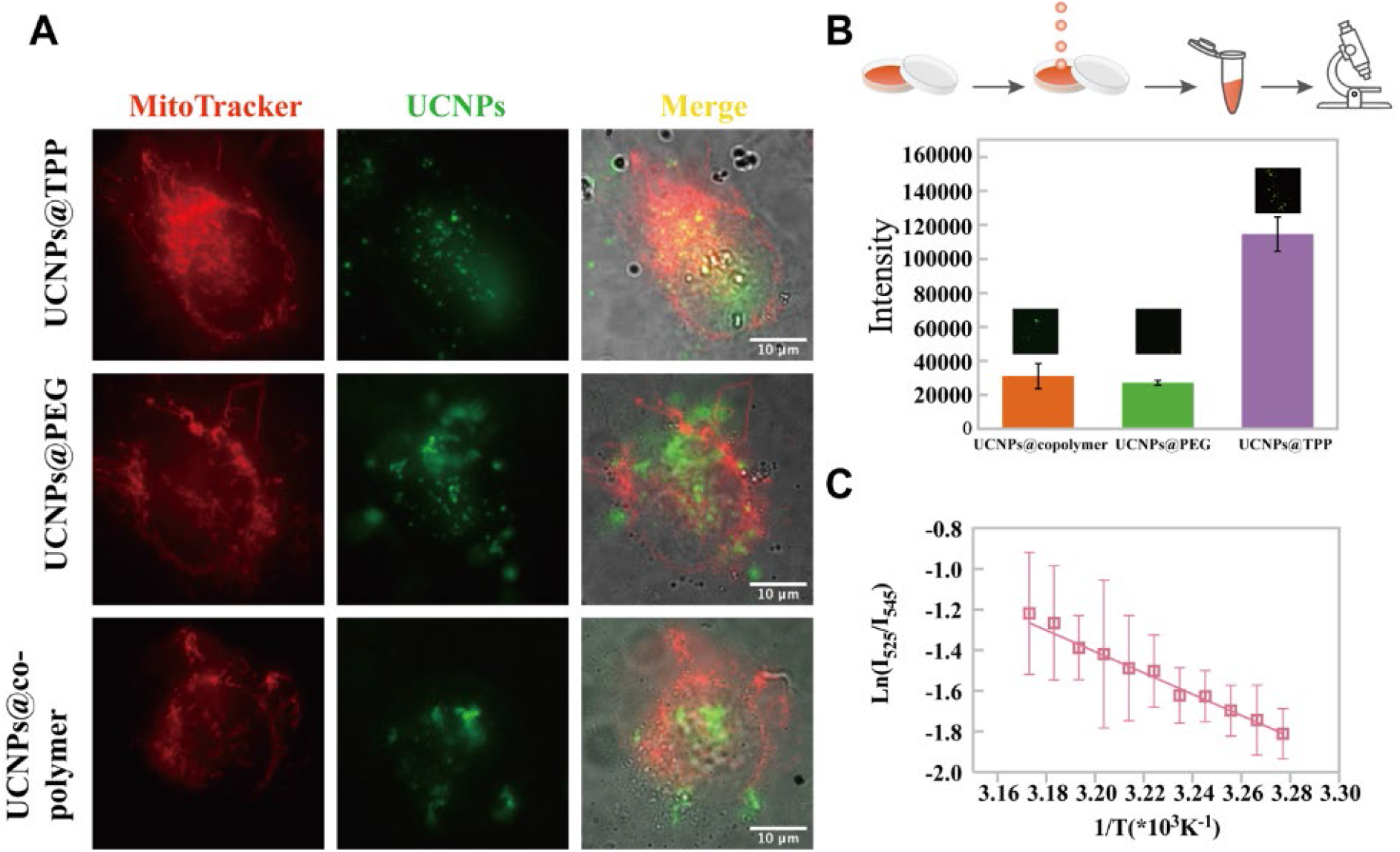
Temperature-dependent fluorescence characteristics of UCNPs@TPP targeted to in situ mitochondria in HeLa cells. A. Intracellular co-localization of UCNPs with MitoTracker (red channel is MitoTracker Deep Red excited by 647 nm laser; the green channel is UCNPs excited by 980 nm laser; the grey channel is the bright-field images). B. The fluorescence intensity of isolated mitochondria treated with UCNPs (n = 10 fields of view from 3 independent repeats). C. Plot of Ln(I_525_/I_545_) vs 1/T to calibrate the thermometric scale in HeLa cells (n = 10 cells). Data points represent mean ± s.d. Scale bar: 10 μm.

Furthermore, to rule out the possible interactions between the UCNPs@TPP and lysosomes, HeLa cells were stained with LysoTracker. After 4 hours’ incubation, HeLa cells were washed with PBS. UCNPs@TPP dispersed into HeLa cells with a speed faster than those in control groups, and there were more nanoparticles delivered to the cells, which suggests the facilitating role of mitochondrial targeting moiety TPP. The merged images showed that UCNPs@TPP already escaped from lysosomes, as a result of the good lipophilicity of TPP (**Figure S4**).

The above results allowed us to proceed to plot the calibration curves in HeLa cells. By changing the extracellular temperature of cell culture using a temperature controllable incubator on top of the microscope system, the logarithmic value of the I_525_/I_545_ ratio showed much more gradual and linear fluorescence changed in the range of 32 to 42 °C relative to the reciprocal temperature (**Figure 3C**), supporting UCNPs@TPP as a quantitative nano-thermosensor in living cells. The equation of calibration curve was y=−5.33x+15.657 (R^2^ = 0.98144, Pearson’s r = −0.99068). The relative sensing sensitivity in HeLa cells at 32 °C is 3.2% K^−1^ and the temperature resolution is ~2.3 K.

### Visualization of mitochondrial thermal dynamics in HeLa cells

We then applied UCNPs@TPP to monitor the mitochondrial temperature variations induced by external nutrient conditions and chemical stimulations. First, we tested a high glucose medium to HeLa cells. Glucose produces pyruvate in the cytosol and then participates in the Krebs Cycle in mitochondria, which generates heat. In the HeLa cells incubated with UCNPs@TPP, the mitochondria temperature increased significantly by 2.25 °C in the first 10 minutes by addition of 5 mg/mL glucose (P < 0.0001 by Students’ t-test) (**Figure 4A**, right), before recovering to the original level after 20 minutes of treatment. As the control group with adding the same amount of PBS, the mitochondrial temperature remained stable within 30 minutes (**Figure 4A**).

**Figure 4.**
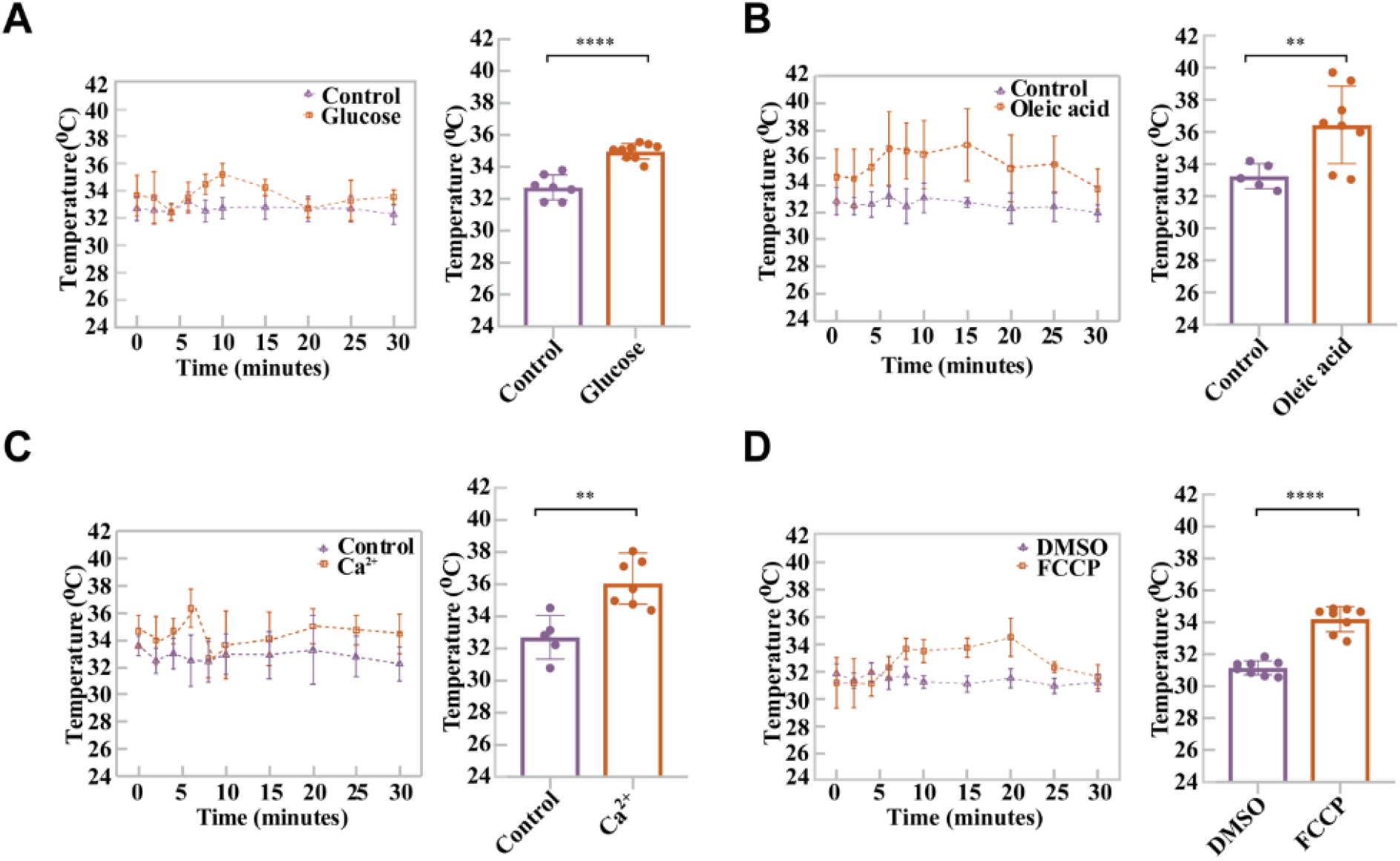
Visualization of mitochondrial thermal dynamics in HeLa cells response to nutrient and chemical stimulations. A. Mitochondrial temperature dynamics in the presence of 5 mg/mL glucose within 30 minutes (left) and Student’s t-test of no glucose and glucose at 10 minutes (p < 0.0001, right). B. Mitochondrial temperature variations in the presence of 5 μM oleic acid within 30 minutes(left) and Student’s t-test of no oleic acid and oleic acid at 15 minutes (p < 0.05, right). C. Mitochondrial temperature changes in the presence of 1 μM ionomycin calcium salt within 30 minutes (left) and Student’s t-test of no calcium and calcium at 6 minutes (p < 0.05, right). D. Mitochondrial temperature fluctuation in the presence of 10 μM FCCP within 30 minutes (left) and Student’s t-test of DMSO and FCCP at 10 minutes (p < 0.0001, right). The temperature in A-D are calculated by the calibration plot in Figure 3C. n = 5~8 cells for A-D, data points represent mean ± s.d.

As an alternative nutrient source, oleic acid was then tested. Columns in **Figure 4B** demonstrates that the mitochondrial temperature increased by 2.74 °C only 5 minutes after adding 5 μM oleic acid in the culture medium, but the mitochondrial temperature didn’t recover to the original level even after 30 minutes of treatment. Compared with glucose, oleic acid treatment makes mitochondrial temperature reach a higher peak value with a faster speed and a longer plateau time, which indicates the different metabolism pathways and energy efficiency for glucose and oleic acid.

The mitochondrial temperature in living cells can be elevated by Ca^2+^ shock as it can promote the pumping of ions and accelerate the respiration reactions ^34^. Ionomycin calcium salt is an ionophore making cell membrane highly permeable of Ca^2+ 35^. Here, **Figure 4C** showed that the temperature of mitochondria increased sharply after adding 1 μM ionomycin calcium salt within 6 minutes before dropping back by another 2 minutes (**Figure 4C**). Ionomycin calcium salt induces intracellular stress, possibly causing damage to mitochondria in HeLa cells ^34^.

Furthermore, carbonyl cyanide-4-(trifluoromethoxy)phenylhydrazone (FCCP) was tested as a chemical stimulation in HeLa cells. FCCP is an inhibitor of oxidative phosphorylation, which disrupts ATP synthesis by transporting protons across the mitochondrial inner membrane ^36^. During this process, mitochondria release a large amount of heat. In the HeLa cells incubated with UCNPs@TPP, the mitochondrial temperature elevated by almost 2 degrees in the first 10 minutes after adding 10 μM FCCP (P < 0.0001 by Students’ t-test) (**Figure 4D**). In the following 20 minutes, the mitochondrial temperature keeps increasing by ~1 °C. Mitochondria eventually recovered to the original temperature after 30 minutes of FCCP treatment. In comparison, the temperature in the control group remained at a similar level after DMSO treatment. These results provide evidence that UCNPs@TPP works as a precise subcellular thermosensor for monitoring the mitochondrial thermodynamics.

## Conclusion and Discussion

Intracellular temperature, especially the mitochondrial thermodynamics, is one of the most crucial biophysical parameters to assess the status of living cells and organisms, which is related to homeostasis and energy balance ^37^. Towards the development of a precise nano-thermosensor both *in vitro* and in living cells, we have synthesized a series of UCNPs@copolymer, UCNPs@PEG, and UCNPs@TPP nanosensors. We applied the non-photobleaching ratiometric nano-thermosensor for monitoring the *in situ* mitochondrial thermodynamics under different physiological and chemical stimuli. The UCNPs@TPP enables us to monitor the glucose-, lipid-, Ca^2+^- and FCCP-dependent thermodynamics in the mitochondria within living HeLa cells. UCNPs@TPP is a powerful tool for analyzing how mitochondria metabolism activates and maintains cellular homeostasis in living cells. The distinct thermodynamics highlight the extensive applications of the thermometer to study vital biological processes related to mitochondrial metabolism and interactions between mitochondria and other organelles, like lysosome, ER ^27^, Golgi ^28^, lipid droplet, and peroxisome ^29^.

Chretien *et al* recently reported that mitochondrial temperature reached >323K (50 °C) using MitoThermo Yellow (MTY) in HEK293T cells treated with an oxygen-rich buffer to fully functionalize the respiration ^38^. Intracellular temperature measurement using organic dyes is not suitable for long-term monitoring purpose. Hu *et al* reported another large increase in temperature by using plasmonic nanostructures with Au nanoparticles in the cytoplasm of CaSki cells during active Ca^2+^ transportation ^39^. Intracellular temperature measurement using inorganic probes requires precise calibration both in the extracellular environment and accurate colocalization of organelles with specific targeting organelle. Fluorescent protein (FP) based thermometer was also reported in HeLa cells by Nakano *et al* with a 6-9 °C temperature increase, which is consistent with our results ^40^. A comparison of different thermometers will be meaningful, including fluorescent proteins ^40^, organic dyes ^13, 38^, plasmonic materials ^39^, UCNPs-based ^23^ nano-thermometer for living cells, etc.

The unique optical properties of UCNPs allow us to track long-term thermodynamics in mitochondria across cell cycles or even in live deep tissues with the NIR excitation wavelength in the future ^10, 11, 41^. The bio-conjugation system we developed will allow us to establish a library of different organelle-targetted nano-thermometers, like lysosome, ER, and Golgi. Combined with other mitochondria evaluation methods, like live-cell super-resolution imaging ^42^, *in vitro* reconstitution assay ^2, 43^ and near-infrared deep tissue imaging ^19^, the UCNPs-based mitochondrial nano-thermometer will be a powerful platform for multifunctional imaging, sensing ^44^, therapy ^19^ and even tracking the pace of life ^45^.

## Data Availability

The data that support the findings of this study are available from the corresponding author on reasonable request.

## Author Contributions

X.D., Q.P.S., J.Z. M.S. and D.J. designed the project; X.D. synthesized UCNPs and performed modification; X.D., D.W., L.Z. and Q.P.S. built optical system, performed experiments and analyzed data; X.D., Q.P.S. and D.J. co-wrote the paper; Q.P.S. and D.J. co-supervised the studies.

## Acknowledgements

This work was supported by the grants from the Australia National Health and Medical Council (NHMRC, APP1177374 to Q.P.S.), the Australia National Heart Foundation (NHF, 102592 to Q.P.S), the University of Technology Sydney’s Grant for IBMD (Q.P.S., X.D., J.Z. and D.J.), the China Scholarship Council (No. 201706170028 to X.D. and No. 201706170027 to D.W.), Australian Research Council (ARC) Future Fellowship Scheme (FT 130100517 to D.J.), and ARC Discovery Early Career Researcher Award Scheme (DE180100669 to J. Z.).

---

**scheme 1.**
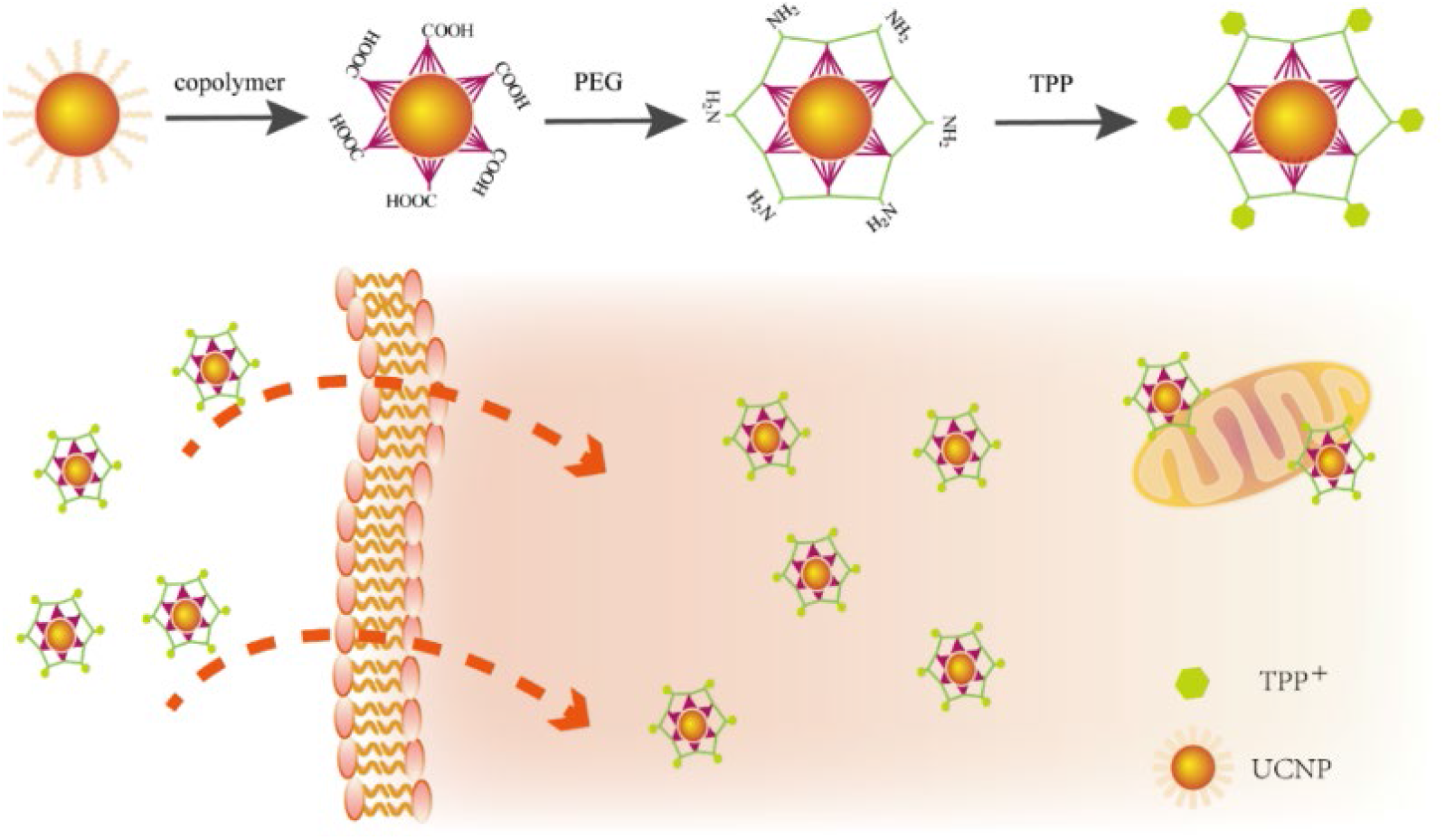
Scheme diagram shows the modification and conjugation of UCNPs and the selective accumulation of UCNPs@TPP to the mitochondria through incubation.

