## Supporting Information for "Quantitatively Monitoring *in situ* Mitochondrial Thermal Dynamics by Upconversion Nanoparticles"

**Support Information for**

**Monitoring *in situ* Thermal Dynamics of Mitochondria by**

**Non-photobleaching Upconversion Nanoparticles**

**Materials and Methods**

**Reagents.** For upconversion nanoparticles (UCNPs) synthesis, yttrium(III) chloride hexahydrate (YCl_3_•6H_2_O, 99.99%), ytterbium chloride hexahydrate (YbCl_3_•6H_2_O, 99.99%), erbium chloride hexahydrate (ErCl_3_•6H_2_O, 99.9%), ethanol, cyclohexane, 1-octadecane, and oleic acid were purchased from Sigma-Aldrich. For surface modification of UCNPs, PEGMEMA_80_-*b*-EGMP_3_ was synthesized by Lin Zhang (Ph.D. candidate in the University of New South Wales from Martina Stenzel group). 4 Arm-PEG-NH2 was purchased from Laysan Bio, Inc. (3-carboxypropyl)triphenylphosphonium bromide (TPP), 1-Ethyl-3-(3-dimethylamino-propyl)-carbodiimide (EDC), N-Hydroxysuccinimide (NHS), MES buffer, HEPES buffer, Tetrahydrofuran (THF), N, N-Dimethylformamide (DMF) were purchased from Sigma-Aldrich with reagent grade or higher. For cell experiments, Dulbecco’s Modified Eagle Medium (DMEM), fetal bovine serum (FBS), Phosphate-Buffered Saline (PBS), Penicillin-Streptomycin (PS), MTT (3-(4,5-Dimethylthiazol-2-yl)-2,5-Diphenyltetrazolium Bromide), MitoTracker (MitoTracker™ Deep Red FM, Invitrogen™ M22426), LysoTracker (LysoTracker™ Deep Red FM, Invitrogen™ L12492), and Mitochondria Isolation Kit were purchased from Life Technologies. Bovine Serum Albumin (BSA), dimethyl sulfoxide (DMSO), glucose, oleic acid-albumin from bovine serum liquid, carbonyl cyanide-4-(trifluoromethoxy)phenylhydrazone (FCCP) and ionomycin calcium salt were purchased from Sigma-Aldrich with reagent grade or higher.

**Synthesis of hydrophobic UCNPs.** NaYF_4_: 20%Yb^3+^, 2%Er^3+^ nanoparticles were synthesized following the previously reported protocols with some modification ^1^. In a typical process, 6 mL of oleic acid and 15 mL of 1-octadecane were added into a 50 mL round bottom flask with three necks, and then added 1.95 mL of YCl_3_ stock solution, 1.0 mL of YbCl_3_ stock solution and 0.2 mL of ErCl_3_ stock solution. The mixture was stirred under argon protection and then heated to 160 °C to get rid of methanol and H_2_O. Second, 5 mL of NaOH-NH_4_F methanol solution was added into the mixture until it cooled down to 30 °C and stirred for 30 minutes. Subsequently, the mixture was heated to 100 °C for 30 minutes and then heated to 300 °C for 1.5 hours. Finally, 5 mL of ethanol was added into the mixture to precipitate UCNPs. UCNPs were washed 3 times with ethanol and cyclohexane before using.

**Modification of UCNPs.** (1) The first step was to modify the UCNPs surface with PEGMEMA_80_-*b*-EGMP_3_ di-block copolymers ^2^. 5 mg of UCNPs and 5 mg of PEGMEMA_80_-*b*-EGMP_3_ di-block copolymers were dissolved in 1 mL of THF and then shaken at room temperature for 12 hours. Next, the UCNPs coated with di-block copolymers (UCNPs@copolymer) were washed with THF and DI water. Finally, the UCNPs@copolymer was dissolved in 0.5 mL of DI water for further use.

(2) The second step was to conjugate UCNPs@copolymer with 4Arm-PEG-NH_2_ ^3^. 5 mg of UCNPs@copolymer, 10 mg of EDC and 10 mg of NHS were dissolved in 1 mL of MES buffer (20 mM, pH 6.0) and shaken for 30 minutes for activation. After centrifugation, the activated products and 10 mg of 4Arm-PEG-NH_2_ were dissolved in 1 mL of pH 7.02 HEPES buffer and reacted overnight. On the second day, reaction products were washed 3 times with DI water. Then nanoparticles were centrifuged at 4400 rpm for 5 minutes to remove the large aggregates. Finally, UCNPs@copolymer conjugated with 4Arm-PEG-NH_2_ (UCNPs@PEG) were stocked in 0.5 mL of DI water.

(3) The third step was to conjugate UCNPs@PEG with TPP. 30 mg of TPP, 10 mg of EDC and 10 mg of NHS were first dissolved into 1mL of DMF by ultrasound. After 1h, 10 mg of UCNPs@PEG were added into the mixture. The reaction was stirred at room temperature for 12 hours and then washed with DMF and DI water. Finally, the product UCNPs@PEG conjugated with TPP (UCNPs@TPP) were stored in 0.5 mL of DI water.

**Characterization.** The morphology of UCNPs was characterized using the FEI Tecnai transmission electron microscopy (FEI, U.S.A.). The hydrodynamic size and zeta potential of UCNPs were determined by a zeta sizer nano (Malvern, U.K.).

**Fluorescence stability of UCNPs@TPP against environmental parameters.** The spectrometer was used to measure the emission spectrum of UCNPs@TPP under different pH, ionic strength (Mg^2+^, Ca^2+^ and K^+^) and refractive index (*n*). The PBS with gradient pH values (4 - 10) were tuned by adding different amounts of HCl or KOH. Ionic strength (0 - 500 mM KCl) was controlled by adding different amounts of KOH. 0 - 2.0 mM CaCl_2,_ and 0 - 1.2 mM MgCl_2_ were obtained by adding different amounts of CaCl_2_ and MgCl_2_. The solutions with different refractive indexes were achieved by glucose. 30% glucose (*n* = 1.3805), 36% (*n* = 1.3912) glucose and 52% (*n* = 1.4222) glucose. 1 mg of UCNPs@TPP were dissolved in the PBS with different pH, different concentrations of ionic strength, or different refractive indexes and transferred to the cuvette. UCNPs@TPP were excited at 980 nm and the spectrum was captured from 480 nm to 600 nm.

**Long-term bio-stability of UCNPs@TPP.** 0.5 mg of UCNPs@TPP were added into 1 mL of DMEM containing 10% FBS or DMEM containing 2% FBS and 0.5% BSA. The hydrodynamic size was applied to monitor the stability of UCNPs@TPP for 7 days.

**Cytotoxicity Assay.** HeLa cells were purchased from ATCC. First of all, 5 mg of MTT was dissolved into 1 mL of sterile PBS buffer. Then 10,000 cells were seeded in the 96-well plate. UCNPs@TPP (0, 1, 10, 50, 250, 500, 1000 µg/mL) were added into each well. Then fresh culture medium was added into the plate after 24 hours. 10 µL of MTT stock solution was added to each well. This 96-well plate was kept in the 37 °C incubator for another 4 hours. Finally, 50 µL of DMSO was added into each well. The microplate reader (Infinite M200 PRO) was applied to read absorbance at 540 nm.

**Colocalization of UCNPs with MitoTracker or LysoTracker.** The HeLa cells were seeded on the fluoro-dish (35 mm) at 10^5^ cell density and incubated in DMEM medium containing 10% v/v FBS for 12 hours. Then the cells were washed 3 times with PBS and incubated with UCNPs (50 µg/mL, 1 mL) for 12 hours (mitochondria colocalization experiment) or 4 hours (lysosomes colocalization experiment) at 37 °C with 5% CO_2_. Next, the cells were washed with PBS and incubated with 200 nM MitoTracker or 200 nM LysoTracker for 0.5 hours. Finally, the cells were washed with PBS and waiting for imaging in DMEM.

**Mitochondria isolation and Fluorescence Intensity Analysis.** Mitochondria from HeLa cells were isolated by a kit bought from Thermo Fisher. Briefly, following the manufacturer’s manual. The protease inhibitors were added into Reagent A and Reagent C before starting the experiment. 2×10^7^ cells were cultured in two flask T175 and treated with 50 μg/mL UCNPs@copolymer, UCNPs@PEG and UCNPs@TPP for 12 hours. Then 2×10^7^ cells were collected by centrifugation. 800 µL of Reagent A was added into the cell pellet and vortexed at medium speed for 5 seconds. The cell suspension was put on the ice for exactly 2 minutes. Next, 10 µL of Reagent B was added in the cell suspension and vortexed at maximum speed for 5 seconds. The mixture was incubated on the ice for another 5 minutes. After 800 µL of Reagent C were added into the cell suspension, the mixture was centrifuged at 700*g for 10 minutes at 4°C. Then the supernatant was transferred to a new tube and centrifuged at 12,000×g for 15 minutes at 4°C. The cell pellet was dissolved in the 500 µL of Reagent C and centrifuged at 12,000×g for 5 minutes. Finally, the pellet was re-suspended in the 50 µL PBS buffer and transferred to a 96-well plate. The fluorescence intensity was recorded by a homemade Total Internal Reflection Fluorescence (TIRF) Microscopy with an excitation wavelength at 980 nm.

**Relative temperature sensing sensitivity and accuracy.** The relative temperature sensing sensitivity (S_R_) and accuracy (σT) are defined as follows ^4^:

$$S_{R}= \frac{dRatio}{dT}\frac{1}{Ratio} (1)$$

$$\sigma T= \frac{\sigma R}{S_{R} \cdot R} (2)$$

where dRatio/dT can be obtained from the slope of the calibration curve represented in Figure 3C and σR is the standard deviation of the ratio. The ratio was obtained from the following equation (**Figure 3C**):

$$Ratio= -5.33T+15.657 (3)$$

**Temperature mapping and temperature sensing of mitochondria.** The HeLa cells were cultured in DMEM containing 10% v/v FBS and 1% v/v Penicillin-Streptomycin at 37 °C with 5% CO_2_. Cells were transferred into the fluoro-dish (35 mm in diameter with No.1 coverglass bottom). Then the cells were incubated in 1 mL culture medium containing 50 µg/mL UCNPs@TPP at 37 °C for 12 hours. The UCNPs@TPP labeled cells were imaged on a home-build total internal reflected fluorescent TIRF microscope by a 980 nm laser excitation under different temperatures (from 30 °C to 42 °C). The homemade TIRF microscopy was installed with an external temperature controller.

For the glucose, oleic acid, FCCP, and Ca^2+^-induced temperature changes, first, the UCNPs@TPP were incubated with HeLa cells for 12 hours. Then, the HeLa cells were stained with MitoTracker. Next, glucose (5 mg/mL), oleic acid (5 µM), FCCP (10 µM) or ionomycin calcium salt (1 µM) was incubated with HeLa cells. The fluorescence intensity ratio (I_525_ / I_545_) of UCNPs@TPP was determined at 980 nm laser.

**Statistical Analysis**

Student’s t-test was applied to examine the differences among variables. Data were shown as mean ± SD. *p values ≤ 0.05 are considered to be statistically significant.

**Expended data**

**Figure. S1**

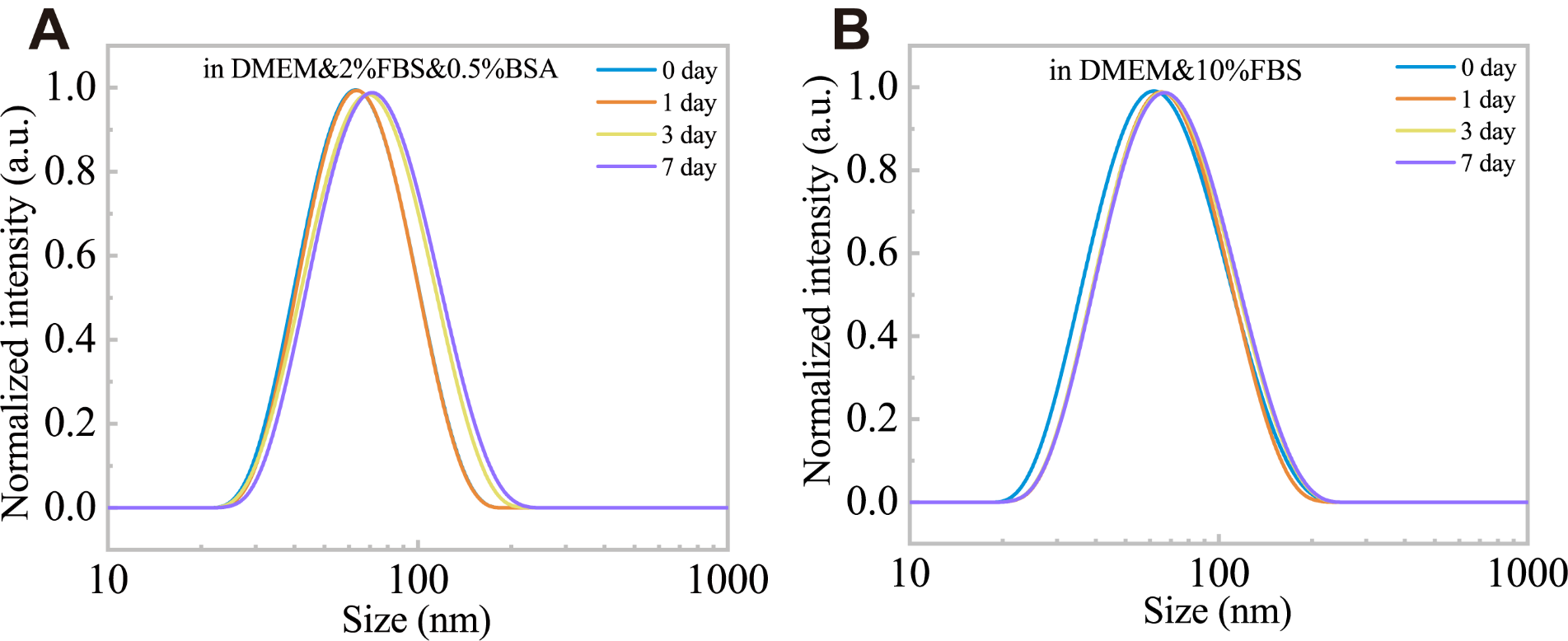

**Fig. S1** Long-term biostability of UCNPs@TPP. A. Bio-stability of UCNPs@TPP in DMEM containing 10%FBS, B. Bio-stability of UCNPs@TPP in DMEM containing 2%FBS and 0.5%BSA. The size of UCNPs@TPP in different physiological buffer keep stable within 7 days.

**Figure. S2**

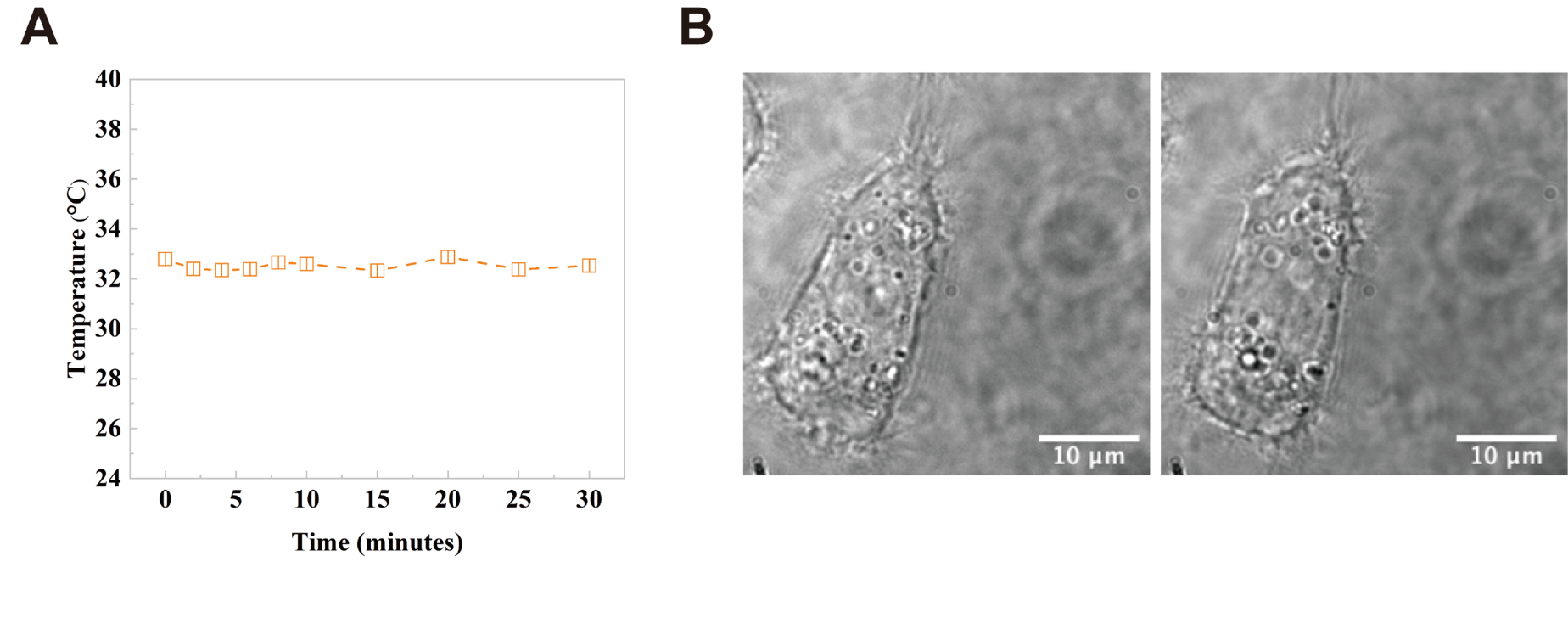

**Fig. S2** The phototoxicity of 980 nm laser. A. The temperature changes of UCNPs@TPP in the water under 0.5 KW/cm^2^ power density and 0.5 seconds exposure time within 30 minutes. B. The shape of the HeLa cell in the 0 and 30 minutes. The power of 0.5 KW/cm^2^ is hardly to elevate the temperature of the water or particles.

**Figure. S3**

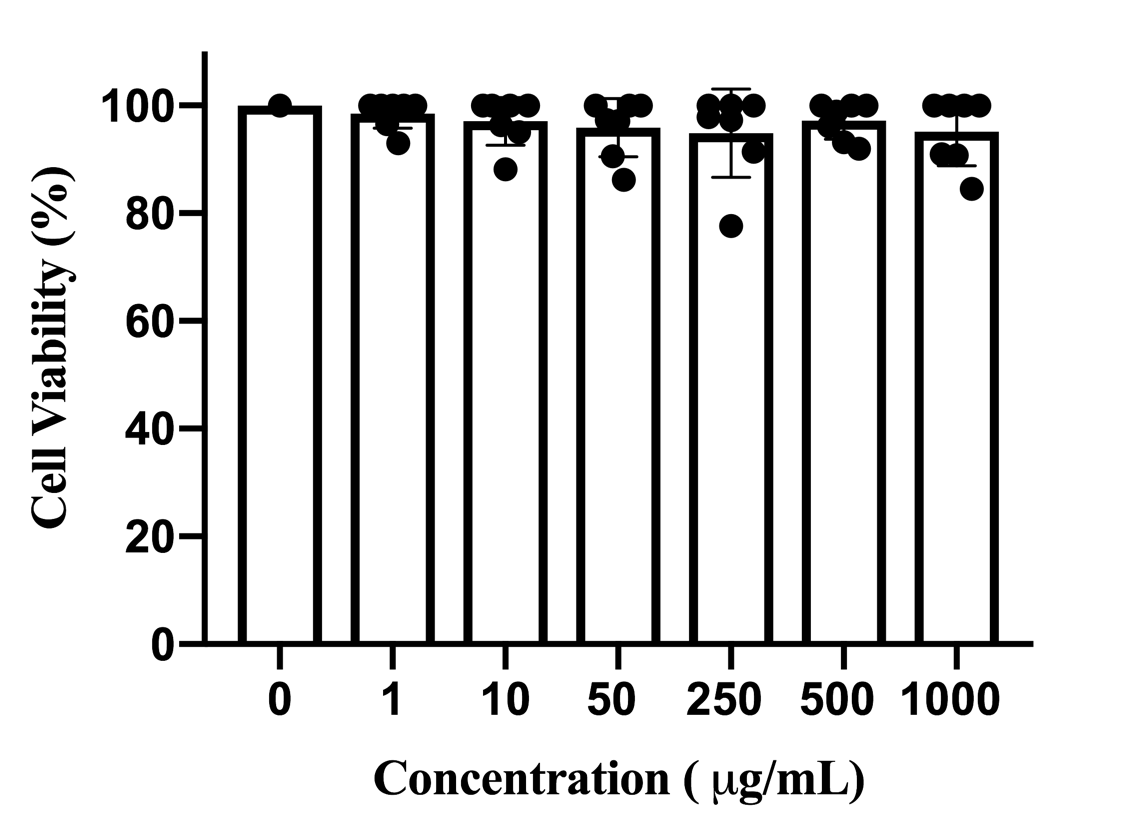

**Fig. S3** The cell toxicity of UCNPs@TPP achieved by the MTT experiment. The concentration of UCNPs@TPP is 0, 1, 10, 50, 250, 500, and 1000 µg/mL. These nanoparticles are low toxicity.

**Figure. S4**

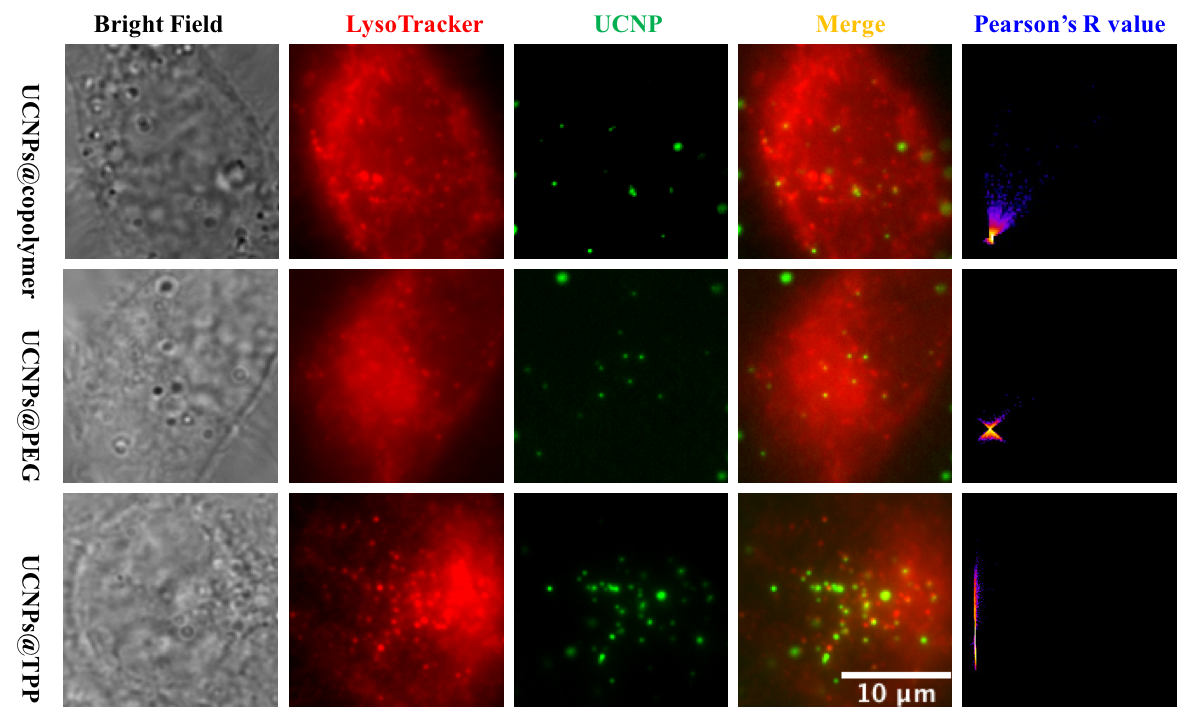

**Fig. S4** Intracellular co-localization of UCNPs with LysoTracker (Red channel is LysoTracker Deep Red excited by 647 nm laser. Green channel was UCNPs excited by 980 nm laser).
